# HDAC11 Regulates RNA Splicing via De-Fatty Acylation of SF3B2

**DOI:** 10.64898/2026.02.22.707301

**Authors:** Jenna Clements, Sung Jung, Ji Cao, Lei Sun, Ana Carolina Ghezzi, Mitchell Gaylen, La’Quan Reid, Changmin Peng

## Abstract

Histone deacetylase 11 (HDAC11) is a lysine de-fatty acylase whose cellular substrates and mechanisms remain incompletely defined. Here, using metabolic labeling, mass spectrometry, click chemistry, and standard molecular biology, we show that SF3B2 is modified by lysine myristoylation at K10 and that HDAC11 efficiently removes this modification in cells, establishing SF3B2 as a direct enzymatic substrate. A de-myristoylation mimetic mutant (SF3B2 K10R) exhibits altered pre-mRNA binding activity in a context-dependent manner. In HCC cells, loss of SF3B2 lysine myristoylation enhances SF3B2 association with androgen receptor (AR) splice variant loci and promotes alternative splicing towards the AR-v7 variant. Consistently, HDAC11 overexpression increases, and HDAC11 knockdown decreases, the AR-v7/AR-FL splice isoform ratio in HCC cells in a manner requiring HDAC11 catalytic activity and recapitulated by SF3B2 K10R. In contrast, modulation of HDAC11 does not alter AR splicing in prostate cancer cells, indicating cell type specific regulation.

Together, these findings establish lysine myristoylation as a reversible regulatory modification on a spliceosomal component and reveal HDAC11-catalyzed de-myristoylation of SF3B2 as a mechanism that can tune alternative splicing in liver cancer cells.

**In Brief:** Clements et al. utilize metabolic labelling, mass spectrometry, click chemistry, and protein and RNA biochemistry to establish that a histone deacetylase enzyme, HDAC11, can influence RNA splicing through de-fatty acylation of the RNA splicing factor SF3B2. De-fatty acylation of SF3B2 at K10 by HDAC11 modulates SF3B2’s pre-mRNA binding to AR splice variant loci, thereby driving alternative splicing of the AR-v7 variant in a cell type dependent manner. This work provides direct mechanistic evidence linking an HDAC to RNA splicing, identifies a reversible lipid modification on SF3B2, and expands current understanding of post-translational regulation of spliceosomal proteins and HDAC11 de-fatty acylation substrates.

**Highlights:**

- HDAC11 de-fatty-acylates SF3B2 at K10, revealing a previously unrecognized modification on SF3B2.
- SF3B2 de-fatty acylation enhances alternative splice-site binding in liver cancer cells.
- HDAC11 regulates RNA splicing through enzymatic de-fatty acylation of a spliceosomal protein.

## Introduction

For decades, the histone deacetylase (HDAC) enzymes have been studied primarily in the context of epigenetics. This has led to major advances in our understanding of gene expression and disease biology^1,2^. Yet HDACs are not only histone deacetylases: many family members efficiently deacetylate non-histone proteins, and several HDACs can remove lysine acylations beyond simple acetylation.^1,3–9^. The intracellular effects of HDAC-mediated deacylation are therefore broad and include protein stability, signaling, localization, and other aspects of cell biology. One area in which HDACs have been suspected to play a role—but without a definitive enzymatic mechanism—is RNA splicing.

The idea that HDACs might be involved in RNA splicing has been present in the field for some time^10^. However, evidence has largely been indirect. Prior work supported a co-transcriptional model in which histone deacetylation alters chromatin accessibility or transcription kinetics, which then correlates with alternative splicing outcomes^11,12^. While these observations are important, they do not establish that an HDAC enzyme can regulate splicing by modifying the splicing machinery itself. One of the more intriguing hints towards HDACs in splicing came from proteomic interactome studies: among HDAC family members, HDAC11 was notable for strong interactions with RNA-associated proteins^13^. It was also observed that HDAC11 knockdown could alter intron retention, but without a catalytic mechanism clearly connecting HDAC11 to RNA biology ^13^.

A major shift in perspectives on HDAC biology came when HDAC11 was discovered to be a potent lysine de-fatty acylase^14–16^. In fact, de-fatty acylation by HDAC11 is >1,000 fold more efficient than de-acetylation^14^. This matters because HDAC11 was cloned in 2003^17^, but its de-fatty acylase activity was only established in 2018–2019^14–16^. As a result, much of the literature on HDAC11—including past work relating to RNA biology—has not been framed through the lens of lysine de-fatty acylation. Given the long-standing suspicion of HDAC involvement in splicing ^10^, the striking RNA-protein interactome profile of HDAC11^13^, and the discovery that HDAC11 is a specialized de-fatty acylase ^14–16^, we hypothesized that HDAC11 could regulate RNA splicing through de-fatty acylation of a splicing regulator.

Here, we report that HDAC11 removes lysine myristoylation (K-myr) from splicing factor 3 subunit 2 (SF3B2), a member of the SF3B complex of the U2 major spliceosome. Using SILAC metabolic labeling, mass spectrometry, and click chemistry, we identify a novel lysine myristoylation site on SF3B2 at K10. We then show that HDAC11 efficiently de-myristoylates SF3B2, establishing SF3B2 as an HDAC11 substrate and linking HDAC11 catalytic activity to a core RNA splicing factor.

SF3B2 has been studied in the context of castration resistant prostate cancer (CRPC), where it promotes alternative splicing of androgen receptor (AR) variant 7 (AR-v7) through binding upstream of variant cryptic exon loci^18^. AR-v7 is well known in CRPC, but AR splice variants are also transcriptionally active oncogenic drivers in hepatocellular carcinoma (HCC), despite low and heterogeneous expression in established HCC cell lines^19,20^. How AR-v7 is regulated in HCC has remained unclear. Thus, AR-v7 provided a defined and biologically relevant splicing readout to test whether HDAC11 de-fatty acylation of SF3B2 could modulate splicing in two cancer types.

We find that loss of SF3B2 K10 myristoylation, modeled by a K10R de-myristoylation mimetic mutant, changes SF3B2 pre-mRNA binding in a cell type–specific way: in HCC cells, de-myristoylated SF3B2 shows enhanced binding at AR-v7 splice variant loci, whereas in prostate cancer cells the effect is not observed as a biologically relevant splicing phenotype. Consistent with this, HDAC11 overexpression and knockdown modulate AR-v7 alternative splicing in HCC cells in a manner that requires HDAC11 catalytic activity and can be phenocopied by the SF3B2 K10R mutant. Together, these data establish a direct enzymatic mechanism connecting HDAC11 to RNA splicing via reversible lysine myristoylation of SF3B2, and they reveal a cancer-type–specific splicing regulatory axis in liver cancer cells.

## Results

### SF3B2 is a substrate of HDAC11 mediated de-fatty acylation

To identify a potential substrate of HDAC11, we began with mass spectrometry (MS). Like previous studies, we combined *in situ* labelling with the fatty acylation probe Alk14 with stable isotope labelling of amino acids in cell culture (SILAC) to identify proteins with increased fatty acylation in HDAC11 knockdown (KD) mouse embryonic fibroblast (MEF) cells ^14,21^. Differentially fatty acylated proteins were cross referenced with available HDAC11 interactome data from pediatric T-cells^13^ and with available HDAC11 SILAC data from HAP1 cells^14^ to yield only five proteins that appeared in all three datasets: SF3B2, CCT3, CCT4, NUP98, and WDR36 (Figure 1A, Table S1).

**Figure 1.**
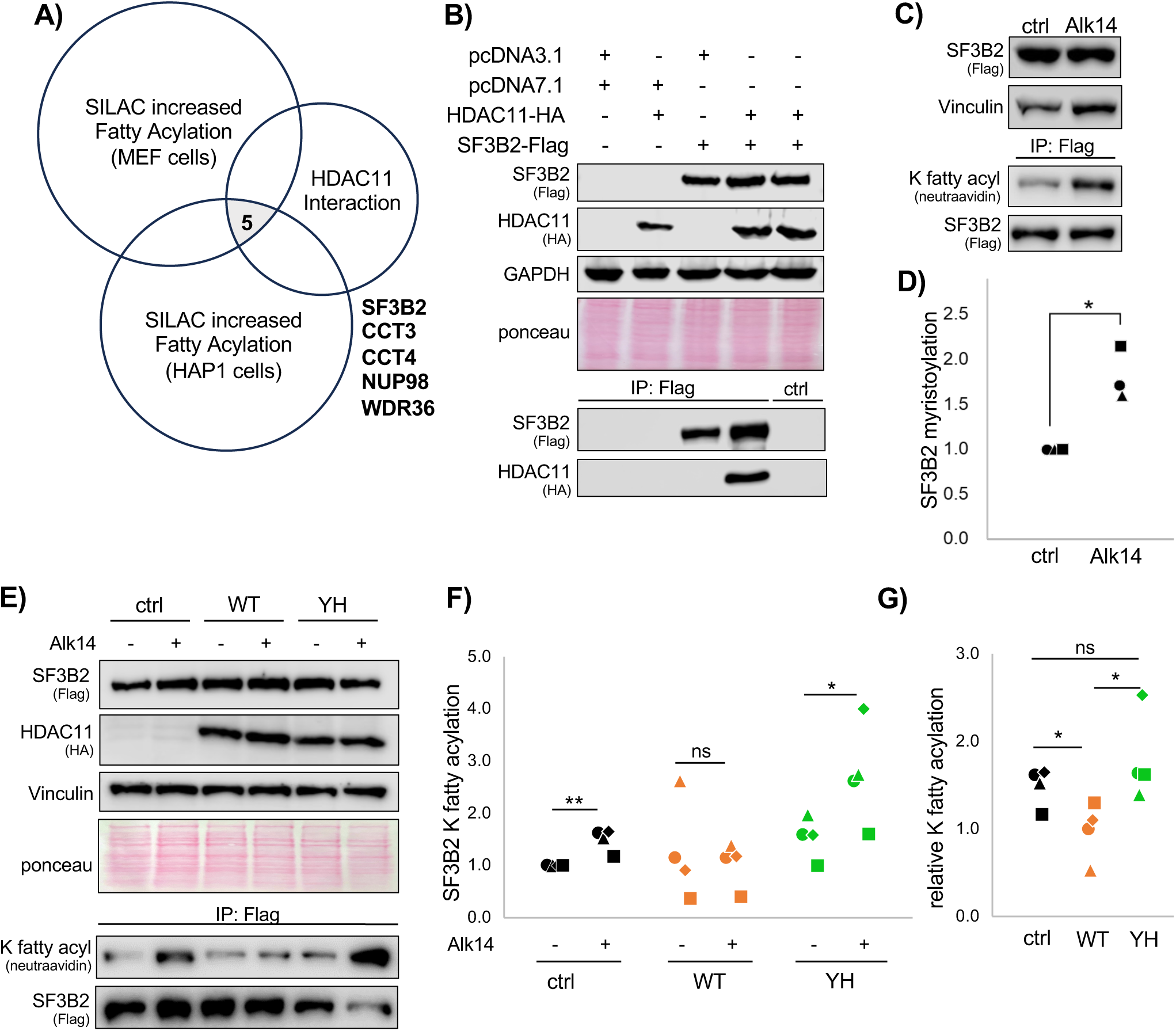
SF3B2 is de-fatty acylated by HDAC11. (**A)** SF3B2 was chosen as a potential substrate after cross-referencing three mass spectrometry datasets. (**B)** Co-IP of SF3B2 and HDAC11. (**C)** SF3B2 lysine fatty acylation by Alk14. (**D)** Quantification of C. (**E)** SF3B2 K fatty acylation with WT or catalytically dead Y304H HDAC11. (**F)** Quantification of E. (**G)** Relative quantification of E. Each condition is normalized by dividing by the +Alk14 signal from its paired control. Each shape in the dot plots represents an independent experiment. Statistics are by paired T-test. *p<0.05, **p<0.01, ns: not significant (p>0.05).

With prior knowledge that HDAC11 is important in HCC^22,23^, we chose SF3B2 for further study because of trends in HCC patient prognosis and expression data that were similar to HDAC11. Both HDAC11 and SF3B2 are individually prognostic towards poor patient outcome in HCC and combining expression of SF3B2 and HDAC11 remained prognostic, indicating correlative expression pattern in the same patients (Figure S1A-C). In HCC patient tumors, both HDAC11 and SF3B2 are significantly increased in the tumor when compared to the normal liver tissue from the same patient (Figure S1D-E). In previous studies, HDAC11 has a direct relationship with HCC cell and tumor growth as well as drug resistance, although the mechanism of this is poorly defined^22,23^. SF3B2, to our knowledge, has not been studied in HCC.

To confirm the interactome data, we tested the HDAC11-SF3B2 interaction in HEK-293T cells. By co-immunopurification (co-IP), HDAC11 and SF3B2 reliably interact (Figure 1B). This interaction is disrupted by loss of function (LOF) mutations that render HDAC11 catalytically dead, pointing towards an enzyme-substrate relationship (Figure S2A). Importantly, HDAC11’s structure has not been obtained, but it is fairly well-predicted by AlphaFold (Figure S2B)^24^. We then used AlphaFold to also predict structures of the HDAC11 mutants, and none of the mutants exhibit a notable structural alteration compared to the WT HDAC11, indicating that the loss of interaction with SF3B2 is due to catalytic function and not structural truncation (Figure S2C-G).

To assay if SF3B2 is a substrate of HDAC11, we used the field-standard method of click-chemistry based readouts with the “clickable” fatty acylation probe Alk14^6,14,21^. SF3B2-Flag was immunopurified (IP’ed) after Alk14 treatment in HEK-293T cells, and biotin was click-conjugated and used for detection by western blot. Reliably, SF3B2 was lysine fatty acylated in HEK-293T cells (Figure 1C-D). Then, SF3B2 fatty acylation was efficiently removed by wild-type, but not Y304H mutant, HDAC11 (Figure 1E-G).

To assay endogenous SF3B2 lysine fatty acylation, we moved to the HCC cell line Hep-G2. For endogenous detection, Alk14 labelled proteins were biotin conjugated prior to streptavidin affinity pulldown of all labelled proteins. Notably, treatment with Alk14 alone seemed to de-stabilized the endogenous SF3B2 (Figure S3A). This may be negated by HDAC11 KD, but overexpression of WT or Y304H mutant HDAC11 did not further alter the SF3B2 protein. We reason the destabilization of SF3B2 is an off-target effect of the harsh nature of the assay. In control cells, SF3B2 was detected in the fatty acylated pulldown fraction, and in HDAC11 KD cells, SF3B2 was increasingly detected relative to the input protein level (Figure S3A-B). This effect was negated by overexpression of WT HDAC11, but not Y304H HDAC11 (Figure S3A-B). This confirms that endogenous SF3B2 can be fatty acylated by Alk14 and de-fatty acylated HDAC11 in Hep-G2 cells.

SF3B2 is nuclear localized, and HDAC11 has been observed in both the nucleus and cytoplasm^17,25,26^. To test HDAC11 localization in Hep-G2 cells, we used basic fractionation of the nucleus and cytoplasm and observed HDAC11 in both compartments to roughly equal levels (Figure S3C). We also tested if knockdown of HDAC11 altered the localization of SF3B2, and it did not; SF3B2 remained concentrated in the nucleus (Figure S3D). Lastly, we tested if HDAC11 displayed histone deacetylase activity in Hep-G2 cells. In a previous study, HDAC11 was suspected to deacetylate Histone 3 at K9 in HCC, but we could not corroborate this result in our Hep-G2 HDAC11 stable KD cell lines (Figure S3E). In sum, data indicates that SF3B2 is a nuclear substrate of HDAC11-mediated de-fatty acylation and that this lipidation of SF3B2 is targeted by HDAC11 more efficiently than Histone 3 acetylation in Hep-G2 cells.

### SF3B2 is myristoylated at Lysine 10

Fatty acylation of SF3B2 is a novel modification, so we moved to validate this by identifying the lysine(s) being Alk14 labelled and de-fatty acylated by HDAC11. We treated HEK-293T cells with fatty acid and IP’ed SF3B2-Flag for detection of lipid post translational modification (PTM) by MS. We found SF3B2 lysine fatty acylation at just one residue, K10, which was lysine myristoylated (Figure 2A). To test this *in situ*, we created a K10R-SF3B2-Flag mutant and found that the K10R construct was no longer fatty acylated by the Alk14 probe (Figure 2B-C). Of note, Alk14 is alkyne palmitic acid, but it can also myristoylate. From this, we concluded that K10 myristoylation (K-myr) must be targeted by HDAC11.

**Figure 2.**
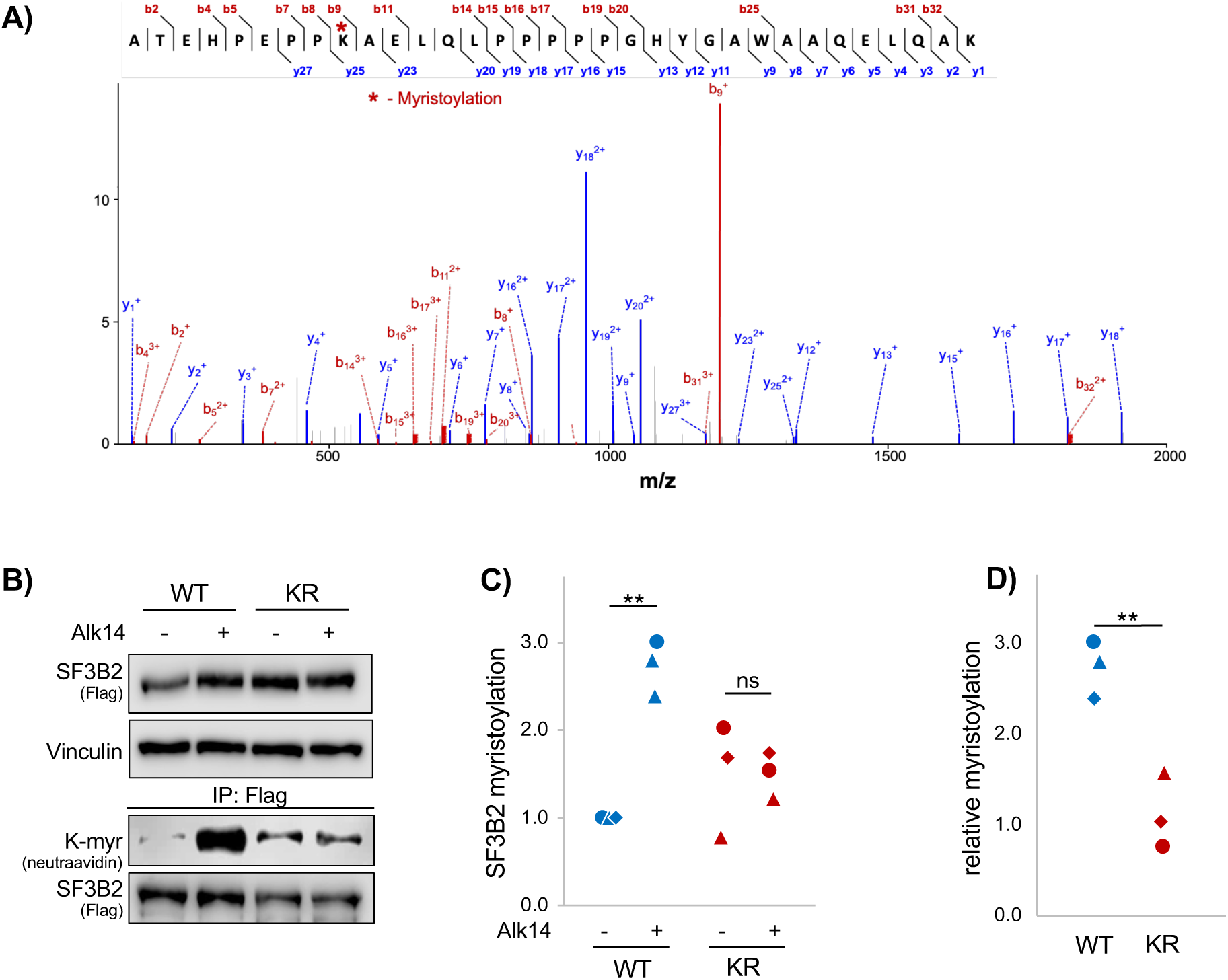
SF3B2 is myristoylated at Lysine 10. **(A)** Mass spectra indicating K-myr at SF3B2 K10. (**B)** Western blot assaying K-myr between SF3B2 and K10R in SF3B2 HEK cells. (**C)** Quantification of B. (**D)** Relative quantification of B. Each condition is normalized by dividing the K-myr signal from its paired control. Each shape in the dot plots represents an independent experiment. Statistics are by paired T-test. **p<0.01, ns: not significant (p>0.05).

There is very little study of SF3B2 post-translational regulation. For example, arginine methylation is necessary for SF3B2’s association with spliceosome machinery and regulates splicing ability^27,28^, and this seems to be the only robustly investigated PTM. To our knowledge, there are no known bona fide SF3B2 lysine PTMs, aside from K-myr. However, SF3B2 is SUMOylated in MS data, and, separately, SUMOylation of spliceosomal proteins is required for efficient pre-mRNA splicing, and broadly effects RNA splicing efficiency^29,30^. There is also some precedent for HDACs modulating lysine PTMs by deacetylation because removal of the acetyl group opens the lysine for other modifications, including ubiquitination and SUMOylation^3,31^. Thus, we hypothesized that de-myristoylation at SF3B2 K10 might modulate SUMOylation at K10.

We expressed SF3B2-Flag with HA-tagged SUMOs and found that SF3B2 could be SUMOylated by SUMO1 (Figure S4A). We used GPS-SUMO to predict the likely SUMOylation sites, and K10 was a top two predicted SUMOylation site (Figure S4B)^32^. Lastly, SF3B2 K10 does constitute a classical SUMOylation motif (Figure S4C). We next tested SUMOylation of the WT SF3B2 versus the K10R SF3B2. Relatively, K10R SUMOylation was lower, indicating that SUMOylation could occur at K10 (Figure S4D). Next, SUMOylation and myristoylation were tested in the same system. SUMO1 expression decreased the K-myr of SF3B2, and conversely, Alk14 treatment decreased SUMOylation (Figure S4E). Additionally, simple overexpression of HDAC11 was sufficient to drastically increase SUMOylation of SF3B2, indicating that HDAC11 may be removing the K-myr to open K10 for SUMO modification (Figure S4F). However, the effects, if any, of HDAC11’s modulation of SF3B2-SUMO are not elucidated by our study. SF3B2 localization and stability, which are two well-known effects of protein SUMOylation^33^, were unaffected by HDAC11 KD (Figure S3B). It is possible that SUMOylation and myristoylation at K10 acts as an on/off switch, and this is further elucidated by direct testing of SF3B2’s pre-mRNA binding function.

### De-myristoylated SF3B2 displays cancer type specific pre-mRNA binding

SF3B2 is one of seven members of the SF3B2 complex, which is part of the U2 complex of the major spliceosome, which is responsible for splicing of 99% of human genes^34,35^. In the context of cancer, SF3B2 has been studied alone in PCa where it was established that SF3B2 regulates Androgen Receptor (AR) alternative splicing^18^.

The AR gene is comprised of eight constitutive exons (Figure 3A). Exon 1 encodes the N-terminal domain, exons 2-3 encode the DNA binding domain, exon 4 codes for the hinge domain, and exons 5-8 make up the ligand binding domain. SF3B2 binds pre-mRNA just upstream of AR cryptic exon 3, located between exons 3 and 4, and promotes its inclusion. When cryptic exon 3 is included, the resulting AR-v7 protein is truncated and it lacks the ligand binding domain. Thus, AR-v7 is ligand-independent and constitutively active^36,37^. AR-v7 is critically important in castration resistant PCa (CRPC) and very well studied in CRPC. In HCC, AR-v7 is emerging as a transcriptionally active oncogene despite low expression levels^19,20^. Yet, in HCC, it is still not known how AR-v7 is regulated. Thus, we moved to test the normal, WT SF3B2 and the K10R de-myristoylated mimic pre-mRNA binding ability in CRPC and HCC cell lines to determine if there is a regulatory function for SF3B2 de-myristoylation.

**Figure 3.**
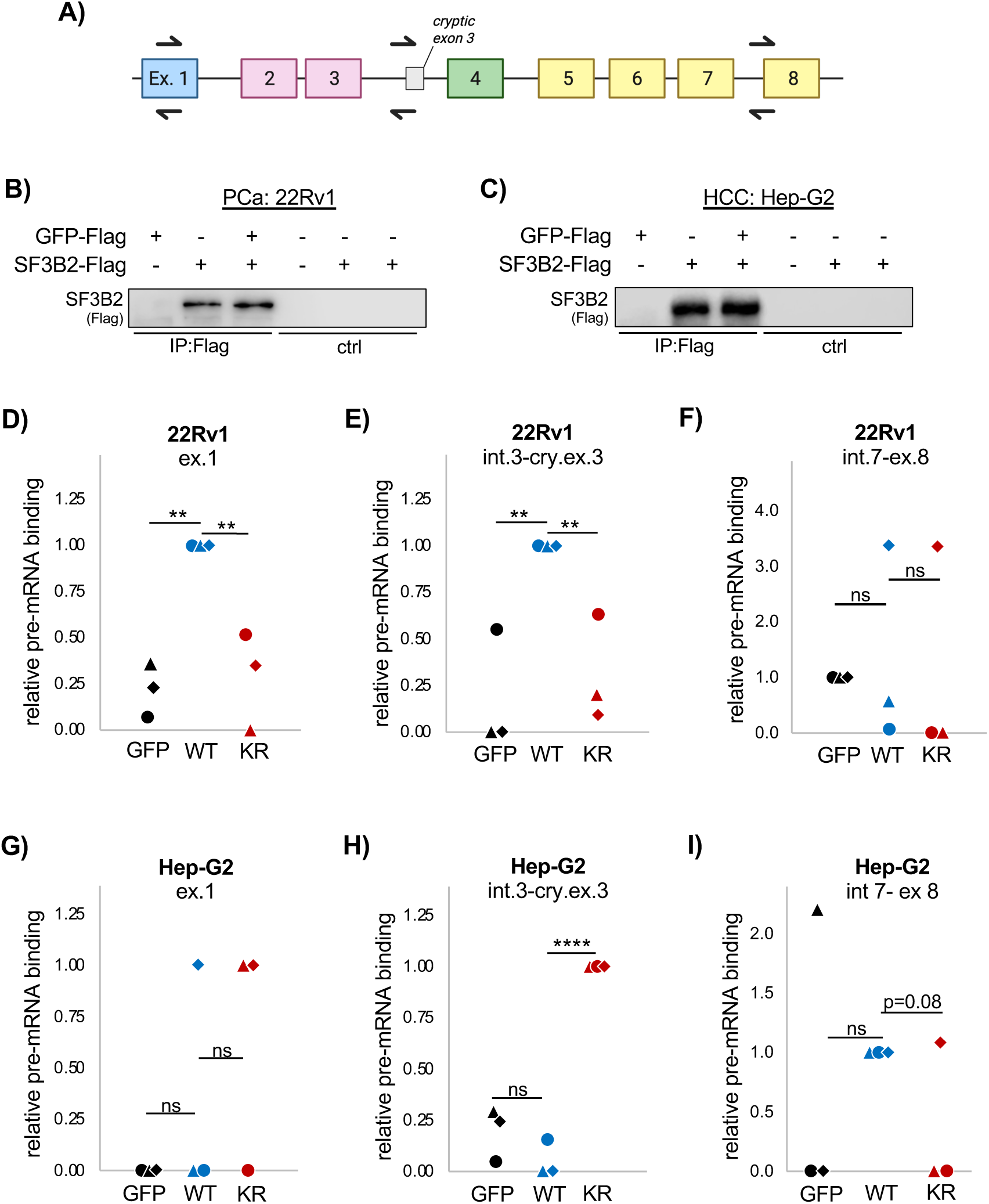
K-myr loss causes cell type specific changes to SF3B2 pre-mRNA binding. **(A)** Schematic of constitutive AR exons, including cryptic exon 3. Exon 1 (blue) encodes the N-terminal domain, exons 2-3 (pink) encodes the DNA binding domain, exon 4 (green) encodes the hinge domain, and exons 5-8 (yellow) encode the ligand binding domain. (**B)** RNA-IP of SF3B2-Flag in 22Rv1 prostate cancer cells. (**C)** RNA-IP of SF3B2-Flag in Hep-G2 hepatocellular carcinoma cells. (**D-F)** SF3B2 WT or K10R binding in 22Rv1 cells at exon 1 (D), intron 3-exon 3 (E), or intron 7-exon 8 (F). (**G-I)** SF3B2 WT or K10R binding in Hep-G2 cells at exon 1 (G), intron 3-exon 3 (H), or intron 7-exon 8 (I). Each shape in the dot plots represents an independent experiment. Statistics are by paired T-test. **p<0.01. ****p<0.0001, ns: not significant (p>0.05).

For CRPC, we chose 22Rv1 cells, which express high levels of AR-v7 and were used previously to establish SF3B2 binding sites^18^. For HCC, Hep-G2 cells were used. We used RNA-Immunopurification (RIP) utilizing WT SF3B2-Flag and K10R SF3B2-Flag. GFP was used as the negative control. PCR primers amplifying established SF3B2 splice sites were used for analysis (Figure 3A), and the IP was consistent in both cell lines (Figure 3B,C). In 22Rv1 cells, WT SF3B2 was bound to exon 1 (Figure 3D) and cryptic exon 3 regions (Figure 3E), recapitulating the previous report in this cell line that these are SF3B2 binding sites^18^. Like that previous study, we also tested the region just upstream of exon 8, and also found that SF3B2 was not bound in that region (Figure 3F). At exon 1 and upstream of the cryptic exon 3, the SF3B2 K10R was not bound in 22Rv1 cells (Figure 3D-E). This indicates that de-myristoylation of SF3B2 might ablate SF3B2’s driving of AR-v7 in CRPC and implies a possible anti AR-v7 effect of HDAC11 in CRPC.

In Hep-G2 cells, the effects of SF3B2 de-fatty acylation were not as easily detected, perhaps due to the lower expression of AR in HCC compared to CRPC^20^. In Hep-G2 cells, WT SF3B2 was not reliably detected at exon 1, but the K10R SF3B2 was enriched in two out of three experiments (Figure 3G). At the cryptic exon 3 locus, the K10R SF3B2 was significantly enriched at this locus compared to both GFP and the WT SF3B2 (Figure 3H), indicating that it is the de-myristoylated SF3B2 that can bind AR-v7 loci in HCC. We also tested the intron 7-exon 8 region, as was done previously, and did not detect statistically significant WT or K10R SF3B2 binding (Figure 3I).

These data point towards a potential cell-type specific function for HDAC11. In PCa cells, results indicate that de-myr of SF3B2 could protect against AR-v7 alternative splicing. This is highly clinically relevant due to the critical importance of AR-v7 in castration resistant prostate cancer (CRPC); 22Rv1 is a CRPC cell line. In HCC cells, on the other hand, de-myr of SF3B2 could drive AR-v7 alternative splicing. Therefore, while our study is HCC focused, both 22Rv1 CRPC cells and Hep-G2 HCC cells were used in subsequent experiments examining the effects of HDAC11 overexpression and knockdown on AR-v7 splicing.

### HDAC11 does not regulate AR-v7 splicing in 22Rv1 CRPC cells

To test if HDAC11 can regulate AR-v7 splicing and expression in CRPC cells, we used both overexpression of HDAC11 and stable knockdown of HDAC11 in 22Rv1 cells. The hypothetical result was that HDAC11 expression would be inversely related to AR-v7. With HDAC11 overexpression, there was not a reliable, statistically significant change to AR-v7 or AR-FL protein (Figure 4A). By quantitative PCR (qPCR) analysis of AR exon junctions, there was a pattern of decreased expression to all the tested AR exon junctions, but this was also short of statistical significance (Figure 4B). Likewise, there was no significant change to the AR-v7/FL ratio, as measured by the ratio of cryptic exon 3-4 junction to exon 7-8 junction, as these are mutually exclusive (Figure 4C). Notably, transient overexpression does not effect all cells, so we also used stable HDAC11 KD to assay if HDAC11 KD in 22Rv1 cells could increase AR-v7 protein.

**Figure 4.**
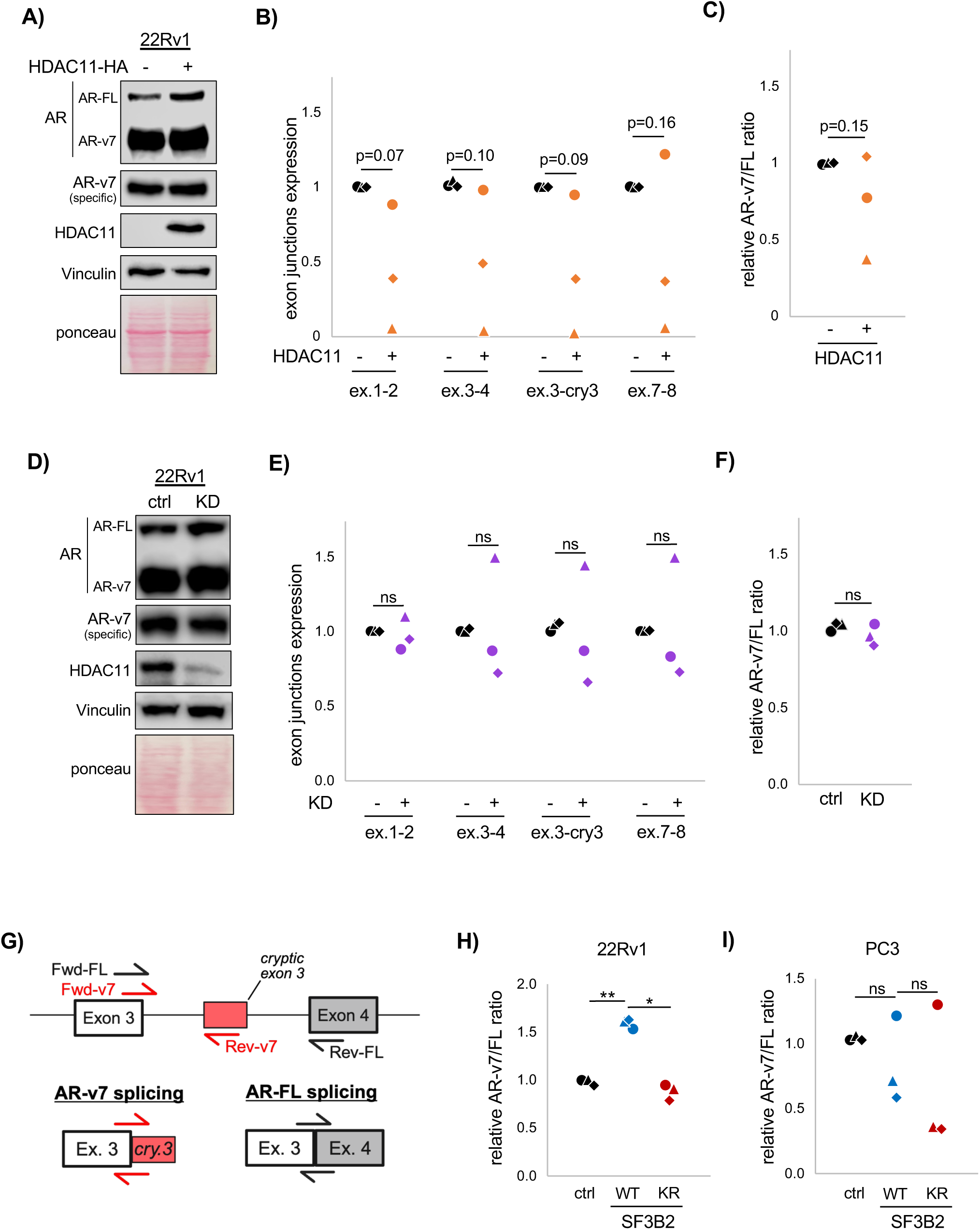
In PCa cells, AR splicing is unaffected by HDAC11 overexpression, HDAC11 KD, or SF3B2 de-myristoylation mimetic. **(A)** Western blot of AR-FL and AR-v7 with HDAC11 overexpression in 22Rv1 prostate cancer cells. (**B)** qPCR analysis of AR exon junctions with overexpressed HDAC11. (**C)** Analysis of AR-v7/AR-FL splicing ratio with overexpressed HDAC11. (**D)** Western blot of AR-FL and AR-v7 with HDAC11 KD in 22Rv1 prostate cancer cells. (**E)** qPCR analysis of AR exon junctions with HDAC11 KD. (**F)** Analysis of AR-v7/AR-FL splicing ratio with HDAC11 KD. (**G)** Diagram of AR-v7 minigene assay. (**H)** AR-v7/FL splicing ratio using minigene in 22Rv1 cells with SF3B2 WT versus K10R de-myr mimetic. **(I)** AR-v7/FL splicing ratio using minigene in PC3 cells with SF3B2 WT versus K10R. Each shape in the dot plots represents an independent experiment. Statistics are by paired T-test. *p<0.05, **p<0.01. ns: not significant.

With HDAC11 KD, there was no affect to AR splicing as measured by western blot or qPCR in 22Rv1 cells. Protein levels of AR-v7 and AR-FL were not altered by HDAC11 KD (Figure 4D), individual AR exon junction expression did not significantly change (Figure 4E), and the ratio of AR-v7/AR-FL was not altered by HDAC11 KD (Figure 4F). Thus, while the RIP data does suggest that SF3B2 de-myristoylation might decrease AR-v7 splicing in CRPC cells, we found HDAC11 manipulation was not biologically relevant to AR-v7 splicing in 22Rv1 CRPC cells.

### HDAC11 regulates AR-v7 splicing through SF3B2 in Hep-G2 HCC Cells

AR-v7 is oncogenic and transcriptionally active in HCC^20^, but the mechanism of its regulation is unknown. A previous report states that established HCC cell lines express barely detectable levels of AR-v7, but despite this, AR-v7 is still transcriptionally active. We compared AR-v7 and AR-FL levels in HCC cells to CRPC 22Rv1 cells and also found that AR-v7 and AR-FL were of very low expression. Neither AR-FL or AR-v7 were detectable by western blot in Hep-G2 or Huh07 cells, while easily detectable in 22Rv1 CRPC cells (Figure S5A). By qPCR, 22Rv1 cells have about 10-fold higher levels of both AR-FL and AR-v7 compared to HCC cells (Figure S5B,C). Thus, in HCC cells, the AR-FL and AR-v7 was still easily detectable, albeit lower than 22Rv1 cells. Lastly, calculation of the endogenous AR-v7/FL ratio within the HCC cells, even compared to the 22Rv1 cells makes clear that while the 22Rv1 cells may express much higher levels of each AR isoform, the ratio of AR-v7/FL in each cell line is comparable. Moving forward, we were able to easily examine the endogenous AR splice patterns after HDAC11 overexpression and knockdown in these HCC cell lines.

In Hep-G2 cells, overexpression of HDAC11-HA (Figure 5A) did not cause significant changes to any of the tested exon junctions when examined alone (Figure 5B). However, with HDAC11 overexpression, paired analysis of the AR-v7/FL ratio across experiments was consistently increased (Figure 5C). The result was similar in Huh-7 cells. HDAC11 overexpression (Figure S6A) did not cause statistically significant changes to AR exon junctions, except for exon 7-8, which was decreased (Figure S6B). But like Hep-G2 cells, the AR-v7/FL ratio was significantly increased (Figure S6C). In these cell lines, the AR minigene assay was applied to assess the effects of de-myr SF3B2 in HCC cells. In both Hep-G2 and Huh-7 cells, the de-myr SF3B2 mimic, K10R, was able to increase AR-v7 splicing (Figure 5D, S6D).

**Figure 5.**
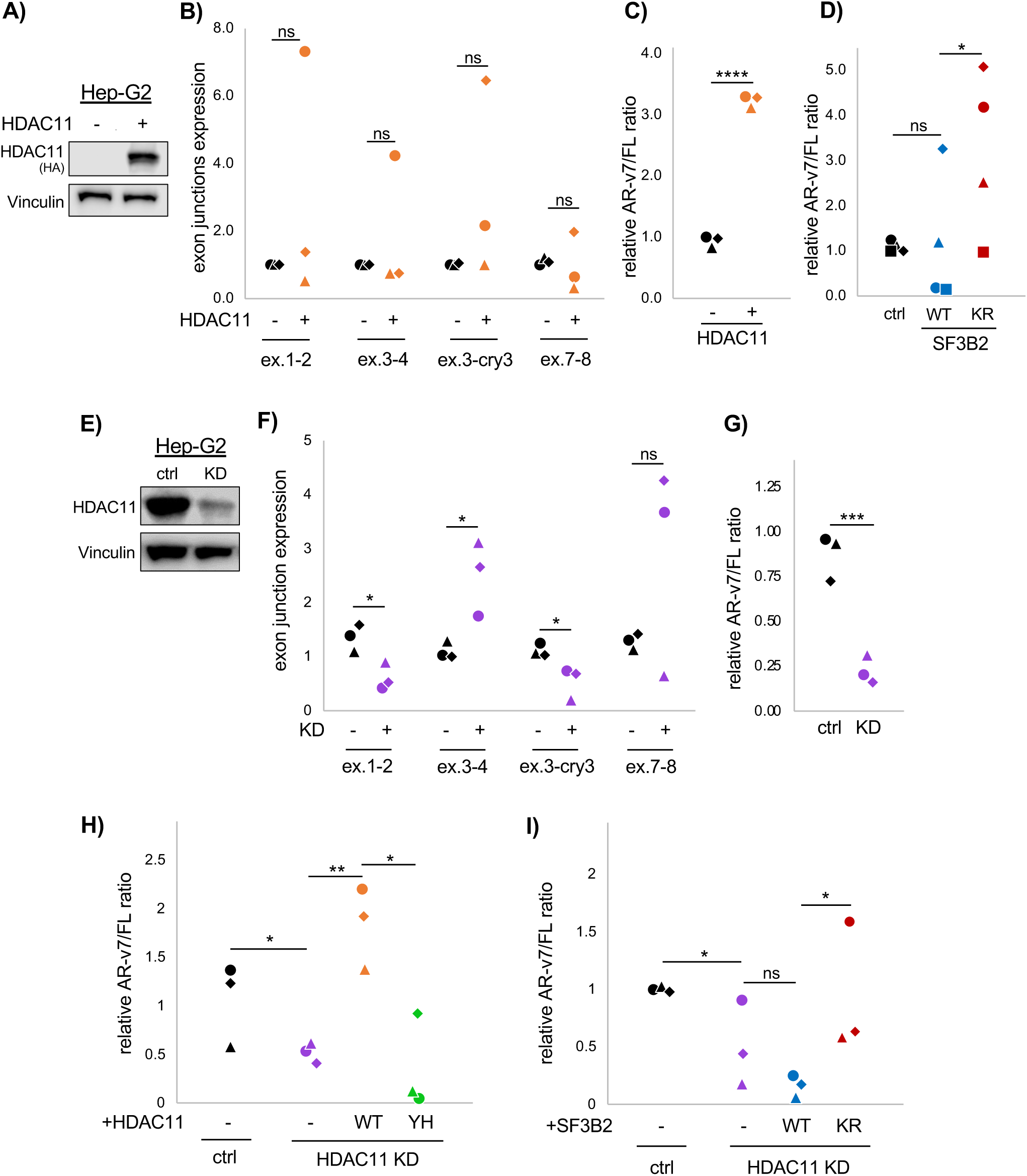
HDAC11 regulates AR-v7 alternative splicing in Hep-G2 cells via SF3B2 de-myristoylation. **(A)** Western blot indicating HDAC11 overexpression in Hep-G2 cells. (**B)** qPCR analysis of AR exon junctions in Hep-G2 cells with overexpressed HDAC11. (**C)** Analysis of AR-v7/AR-FL splicing ratio in Hep-G2 cells with overexpressed HDAC11. (**D)** AR-v7/FL splicing ratio using minigene in Hep-G2 cells with SF3B2 WT versus K10R. (**E)** Western blot indicating HDAC11 KD in Hep-G2 cells. (**F)** qPCR analysis of AR exon junctions in Hep-G2 HDAC11 KD cells. (**G)** Analysis of AR-v7/AR-FL splicing ratio in Hep-G2 HDAC11 KD cells. (**H)** AR-v7/FL splicing ratio in Hep-G2 HDAC11 KD cells with overexpressed WT or mutant HDAC11. (**I)** AR-v7/FL splicing ration in Hep-G2 HDAC11 KD cells with overexpressed WT or K10R SF3B2. Each shape in the dot plots represents an independent experiment. Statistics are by paired T-test. *p<0.05, **p<0.01, ***p<0.001, ****p<0.0001, ns: not significant (p>0.05).

We then tested stable HDAC11 KD in each cell line. In Huh-7 cells, HDAC11 KD did not cause statistically significant changes to individual exon junctions but did decrease the AR-v7/FL ratio, albeit with some variability (Figure S6E-G). In Hep-G2 HDAC11 KD cells (Figure 5E), HDAC11 KD caused a consistent decrease to the exon 1-2 junction and the exon 3-cryptic exon 3 junction (Figure 5F). Exon 1-2 is not specific to AR-v7, and its decrease indicates a possible downregulation to many isoforms of AR. The cryptic exon 3 is specific to AR-v7, however, and its decrease alone indicates decreased AR-v7. There was also a significant increase to exon 3-4 junction, which is in AR-FL and not AR-v7, as well as an increase that was short of statistical significance to exon 7-8, which is in AR-FL and not AR-v7 (Figure 5F). With HDAC11 KD, the ratio of AR-v7/FL decreased significantly, by about 75% (Figure 5G), indicating that HDAC11 KD causes the AR-v7 to AR-FL splice isoform switch in Hep-G2 cells.

We then tested if the decreased AR-v7/FL splice ratio in Hep-G2 HDAC11 KD cells could be rescued by overexpression of WT HDAC11 or Y304H catalytic mutant HDAC11. Only the WT HDAC11 was able to rescue the AR-v7/FL ratio (Figure 5H). This confirms that the splice switch is controlled by HDAC11’s de-fatty acylation function. Next, we also tested if the decreased AR-v7/FL ratio could be rescued by overexpression of SF3B2 K10R, which mimics de-fatty acylation. Indeed, overexpression of K10R SF3B2, but not WT SF3B2, increased the endogenous AR-v7/FL splice ratio (Figure 5I). This result confirms that de-fatty acylation of SF3B2 exists downstream of HDAC11 and works to regulate AR-v7 splicing in HCC cells.

## Discussion

Prior to this study, HDACs involvement in splicing was thought to occur co-transcriptionally. Our work is separated from this model because we demonstrate that HDAC11 directly modifies an RNA splicing factor to influence that splicing factor’s function on nascent RNA. We identify SF3B2 as a de-myristoylation substrate of HDAC11, which regulates SF3B2’s pre-mRNA binding function specifically in HCC cells, and establishes that de-myristoylation of SF3B2 by HDAC11 can regulate AR-v7 alternative splicing in HCC cell lines. 22Rv1 CRPC cells did not have an AR-v7 splicing phenotype downstream of HDAC11. These findings establish HDAC11 as a regulator of RNA splicing and are the first to directly implicate any HDAC family member in the process of splicing itself. Our work also adds to knowledge of SF3B2 regulation, of which very little is presently known, and establishes the first mechanism of AR-v7 splicing regulation in HCC. Separately, we also demonstrated that HDAC11 KD did not alter the previously reported histone target H3K9, supporting a possibility that RNA splicing is HDAC11’s mechanism of gene expression regulation.

The potential effects of HDAC11’s regulation of SF3B2 is vast. SF3B2 is part of the major spliceosome complex, which is responsible for splicing 99% of human genes ^34,35^. We showed that de-myristoylated SF3B2 has enhanced binding at AR-v7 loci, but other potential loci are untested by our study. Other genes influenced by HDAC11, such as LKB1 and Ataxin 10, have known splice isoforms^13,23,38^. It is possible that HDAC11’s regulation of these genes occurs at the post-transcriptional step of RNA splicing but was previously attributed to the transcriptional level due to HDAC11 not being yet well-studied as a de-fatty acylase, let alone studied in splicing itself. Further studies would benefit from deep RNA sequencing to identify splice isoforms under HDAC11 influence; we limited our study to enzymatic mechanism and a basic splicing readout.

Myristoylation of SF3B2 at K10, and de-myristoylation of SF3B2 by HDAC11, is a novel modification and SF3B2 is a novel HDAC11 substrate. Presently, it is not possible to directly test endogenous fatty acylation, that is, without exogenous fatty acid probe treatment, due to a lack of antibodies for these highly hydrophobic, long lipid modifications. Although, there is promise that this can be done in future studies since one group was able to create a palmitoylation antibody specific to TEAD palmitoylation^39^. Previously, understanding of post translational regulation of SF3B2 was limited to just R508 methylation by PRMT9, where loss of R508me causes aberrant RNA splicing and abnormal development^27,28^. Our study also presents SUMOylation as a possible PTM on SF3B2. When SUMO1 was overexpressed, SF3B2 was SUMOylated. This was also modulated by both HDAC11 and Alk14 at K10, but we did not discern whether this is biologically relevant under endogenous conditions because there was no change to SF3B2 localization or stability, which are primary functions of SUMO, with HDAC11 KD. Therefore, it is possible that the cell-type dependent effects observed in our study are caused also by SUMO dysregulation.

The highlight of our study is the effect of SF3B2 de-myristoylation by HDAC11: splice isoform switching of AR-FL to AR-v7 specifically in HCC cells. HCC displays a marked sex dimorphism, and the same is true for fatty liver disease, a leading cause of HCC in the United States^40,41^. There is suspicion that this is due to AR expression in the liver, and other studies have established the oncogenicity of AR and its transcriptional activity in HCC despite low expression^20^. Our study is the first to propose a regulatory mechanism for AR-v7 in HCC, and it is sensible that HDAC11, through SF3B2, both of which are prognostic in HCC, upregulates an oncogene via alternative RNA splicing.

## Materials and Methods

### Cell Culture

All established cell lines were cultured in DMEM, (Cytiva SH30243.FS) except for 22Rv1 cells, which were cultured in RPMI-1640 Medium (Cytiva SH30027.FS). Each medium was supplemented to a final concentration of 10% fetal bovine serum (Cytiva SH30919.03) and 1% penicillin-streptomycin (Cytiva SV30010). All cells were adherent and maintained in 10 cm polystyrene treated tissue culture treated dishes (Fablab #FL7621) and incubated at 37 °C and 5% CO_2_. Cells were passaged when they reached 90-100% confluency; the frequency of passaging varied between cell lines.

### Plasmid construction

SF3B2 and HDAC11 coding DNA (cDNA) was obtained from Sino Biological (HG16923-U and HG11490-M). The cDNAs were inserted into pcDNA7.1-3XFlag or pcDNA3.1-HA vectors by PCR and restriction enzyme digest. Plasmid sequences were verified using whole plasmid sequencing from Plasmidsaurus. To generate the HDAC11 catalytic mutants, D181A, H183A, and Y304H, and the SF3B2 K10R construct, PCR primers targeting the residues of interest were used, and the entire plasmids were amplified with KOD Hot Start Polymerase. All primers are listed in Table S2.

### Transfection and stable cell line selection

For transient transfection, 4e^6^ cells were treated overnight with 18 µg total plasmid DNA per 10 cm plate. For experiments where multiple plasmids were used, the total DNA did not exceed 18 µg. Lipofectamine 2000 (Thermo 11668027) or polyethylenimine (PEI, Sigma 408727) transfection reagents were used at a ratio of 1-3 ug per ug of DNA. DNA and transfection regent were diluted in Opti-MEM^TM^ Reduced Serum Media (Thermo 31985070) before adding to cells in complete medium. The medium was refreshed the next day, and cells were harvested 2 days later.

To generate stable cell lines, lentivirus was generated by co-transfection of shRNA vector targeting HDAC11, psPAX.2, and pMD2.g into HEK-293T cells. The HDAC11 shRNA construct in PLKO.1 vector was generated previously^42^. Virus-containing medium was collected after 48 hours and used to transduce cells with polybrene. Cells were selected with puromycin, and then grown in half the lethal dose of puromycin once selected. Cells were selected for at least 4 days before validating the KD with western blot and beginning experiments.

### SILAC mass spectrometry

SILAC MS was completed as previously described using HDAC11 WT and KD MEF cells^14^. MEFs were isolated and HDAC11 KD took place as previously described^43,44^. In brief, MEFs were isolated from C57BL/6 E14.5 embryos using enzymatic digestion and maintained in DMEM/Hi glucose (HyClone) supplemented with 10% FBS (Gemini), 1% penicillin-streptomycin (Gibco), and 1.1% GlutaMAX (Gibco). Knockdown of HDAC11 was completed with lentivirus containing shRNA targeting HDAC11^43^.

### Detection of SF3B2 myristoylation by mass spectrometry

HEK-293T cells were passaged three times over 10 days in complete growth medium supplemented with high concentration fatty acid. The fatty acids were a 1:1 molar ratio of myristic and palmitic acid: 25 µM myristic acid and 25 µM palmitic acid, which were diluted into the cell culture at 1:1000 from a 100% ethanol vehicle. Cells were transfected with SF3B2-Flag in OPTI-Mem media for 6 hours; the fatty acids were removed during this time. Afterwards, the media was replaced with complete DMEM culture medium supplemented with the high concentration fatty acids. Cells were lysed two days later in denaturing lysis buffer (1% NP-40, 1% SDS, 50 mM Tris, 150 mM NaCl, 10% Glycerol, in PBS with protease inhibitors). IP was completed overnight at 4 °C using 80 mg of total protein divided among 12 mL of lysis buffer in 12 test tubes. The lysis buffer was supplemented to 350 mM NaCl for IP. Each IP aliquot contained 20 µL of Anti-Flag G1 affinity resin (GenScript L00432-5). Resins were washed 3X in NP-40 Wash Buffer (0.05% NP-40, 50 mM Tris, 350 mM NaCl, 10% Glycerol, in PBS) and 2X in PBS. PBS was removed completely needle aspiration and samples were kept at -80 °C until mass spectrometry.

Detection of myristoylation on purified SF3B2 was performed as previously described, with minor modifications^45^. Briefly, affinity-purified proteins were digested on beads using Trypsin/Lys-C. The resulting peptides were enriched and desalted using in-house STAGE tips^46^ packed with 2 mg of C18 beads (3 μm, Dr. Maisch GmbH, Germany), followed by vacuum drying. Dried peptides were resuspended and analyzed on an Easy-nLC 1200 system coupled to an Orbitrap Fusion mass spectrometer (Thermo Scientific) equipped with an electrospray ionization (ESI) source. Data were acquired in data-dependent acquisition (DDA) mode to enable unbiased peptide detection. Raw spectra were processed using Proteome Discoverer (v2.5; Thermo Fisher Scientific) with the Mascot search engine (v2.4; Matrix Science). Dynamic modifications included myristoylation on N-terminal glycine, lysine, and cysteine residues, as well as oxidation of methionine. Peptide identifications were filtered to a 1% false discovery rate (FDR) using Percolator validation based on q-values.

### Detection of SF3B2-HDAC11 interaction

To test protein-protein interaction of HDAC11 and SF3B2, plasmids were transfected at a 1:1 ratio and co-IP was used. HEK-293T cells were lysed in denaturing lysis buffer, probe sonicated, and centrifuged at high speed to clarify the lysate. Samples were brought to equal concentration and volume for co-IP; no less than 2 mg total protein was used in a total volume of 1 mL lysis buffer supplemented to 500 mM NaCl. Anti-Flag G1 affinity resin was used or just Protein G Agarose resin (Thermo 15920010) was used as a negative control. IP took place overnight at 4 °C. The next day, resin was washed 3X in cold NP-40 wash buffer supplemented to 500 mM NaCl. Protein was eluted directly into 1X Gel Loading Buffer and resolved by SDS-PAGE on 7.5% polyacrylamide gels. Western blot proceeded by standard method.

### Detection of SF3B2 lysine myristoylation by western blot in HEK-293T cells

SF3B2-Flag was transfected into HEK-293T cells. Two days later, cells were treated with Alk14 (CCT-1165) 50 µM or DMSO vehicle for 6 hours. Cells were lysed in denaturing lysis buffer, probe sonicated, and centrifuged to clarify the lysate. Lysates were brought to equal concentration and volume before immunopurification (IP). SF3B2 was IP’ed using 20 µL of Anti-Flag G1 affinity resin overnight at 4 °C. The resin was washed 3X in NP-40 wash buffer, followed by the Click reaction to conjugate biotin to Alk14-labelled proteins.

The Click reaction took place with 60 µL of NP-40 wash buffer, 5 µL of 2 mM Biotin Azide (Vector Labs CCT-1265) in DMSO, 5 µL of 100 mM THPTA (Vector Labs CCT-1010) in DMSO, 5 µL of 100 mM sodium ascorbate (Sigma A4034) in water, and 5 µL of 40 mM CuSO_4_ (Sigma C1297) in water. The reaction took place in a thermomixer at 28 °C, 400 rpm, for 90 minutes. Afterwards, the resin was washed in 1 mL NP-40 wash buffer. The wash solution was removed before elution into 50 uL of 4% SDS, pH 8.5, by heating at 95 °C for 10 minutes. Before SDS-PAGE, an aliquot of sample was supplemented with hydroxylamine (Sigma 159417) to final concentration of 500 mM to remove possible cysteine palmitoylation. Samples were heated at 95 °C for 5 minutes, then 6X Gel Loading Buffer was added so that the final concentration of gel loading buffer was 2X. Samples were resolved using 7% polyacrylamide gels and transferred to nitrocellulose membranes for standard western blot analysis. Western blot was completed using NeutrAvidin-HRP (Thermo 13030) diluted at 1:5000 into 5% bovine serum albumin (BSA). Western blot for total Flag was completed with Flag antibody (Table S3) at 1:3000 in 5% BSA.

### Detection of endogenous SF3B2 myristoylation in Hep-G2 cells

Per condition, empty HA vector control, WT HDAC11-HA or Y304H HDAC11-HA was transfected into 10e^6^ Hep-G2 shGFP control or HDAC11 KD cells. Two days later, cells were treated with Alk14 (50uM) or DMSO vehicle control for 6 hours. Cells were lysed in 200 uL of denaturing lysis buffer, probe sonicated, and centrifuged to clarify the lysate. Lysates were brought to equal concentration in NP-40 wash buffer. A small aliquot from each sample was stored at this point for input control analysis. 1 mg total lysate was brought to volume of 800 uL in NP-40 wash buffer. The reaction components were added to final volume of 1 mL: 50 µL of 2 mM Biotin Azide in DMSO, 50 µL of 100 mM THPTA in DMSO, 50 µL of 100 mM sodium ascorbate in water, and 50 µL of 40 mM CuSO_4_ in water. The reaction took place in a thermomixer at 28 °C, 400 rpm, for 90 minutes. Afterwards, protein was extracted.

To extract protein, samples were transferred to 15 mL conical tubes and 4 mL of ice-cold methanol was added followed by vortexing. 1 mL chloroform was added, samples were briefly vortexed, and 3 mL ice cold water was added. Samples were vortexed for 30 seconds and then centrifuged at 14,000 xg. The top aqueous layer was removed, 4 mL methanol was added, and samples were centrifuged again for 5 minutes at 20,000 xg. Methanol was removed, protein was dried, and then protein was resolubilized in 100 uL of 4% SDS pH 8.5 with warming and mild sonication as necessary. Protein solutions were then diluted to 1.5 mL using NP-40 wash buffer, and 20 uL of streptavidin magnetic beads (NEB S1420S) were added to each sample. Biotin-streptavidin affinity purification to capture Alk14 labelled proteins took place at 4 °C for 6 hours with samples turning end-over-end consistently. Beads were separated in a magnetic rack for 2 minutes and washed 3X in 1 mL of ice-cold NP-40 wash buffer. Then, beads were incubated in 0.5 M hydroxylamine for 1 hour to remove potential cysteine acylation by Alk14. Hydroxylamine was removed, beads were washed once more, and then were resuspended in 50 uL of 4% SDS, pH 8.5 and heated at 95 °C for 10 minutes to elute lysine fatty acylated proteins. Samples were resolved using 7% polyacrylamide gels and transferred to nitrocellulose membranes for standard western blot analysis with SF3B2 antibody.

### Analysis of HDAC11 in HCC patient data

Kaplan-Meier plots were generated using the Pan-cancer RNA-seq analysis tool from kmplot.com^47^. Analysis of HDAC11 and SF3B2 expression in normal and tumor samples from HCC patients was generated by tnmplot.com^48^. Both tools pull from TCGA patient data.

### Analysis of HDAC11 Predicted Structures

To predict the structure of HDAC11, the amino acid sequence obtained from sequencing the HDAC11-HA plasmid was input into AlphaFold 3 and the resulting structure was downloaded^24^. The same was done for amino acid sequences of HDAC11 mutants D181A, H183A, and Y304H. To compare the structures of WT HDAC11 to the HDAC11 mutants, the paired structures were analyzed in UCSF Chimera X using Matchmaker and colored by per residue root mean square deviation (RMSD)^49^.

### SUMOylation assays

To assay SUMOylation on SF3B2, co-transfection of SF3B2-Flag and SUMO-HA plasmids was used. The plasmids were transfected at a 2:1 ratio of SF3B2:SUMOs. In experiments where HDAC11 was co-transfected with SF3B2 and SUMO1, the transfection ratio was 3:2:1 SF3B2:HDAC11:SUMO1. In experiments where Alk14 was assayed against SUMOylation, cells were treated with 50 µM Alk14 or DMSO for 6 hours right before lysis. Cells were lysed in denaturing lysis buffer supplemented to 500 mM NaCl, probe sonicated, and centrifuged to clarify the lysate. SF3B2-Flag was IP’ed using 5 mg total lysate in 1 mL lysis buffer with 20 µL Flag affinity resin overnight at 4 °C. Resin was washed 3x in NP-40 wash buffer supplemented to 500 mM NaCl, and protein was eluted into 1X Gel Loading Buffer by heating at 95 °C for 10 minutes. Samples were resolved on 7% polyacrylamide gels and western blotting took place with Flag and HA antibodies according to standard approach.

### RNA Immunopurification (RIP)

22Rv1 or Hep-G2 cells were grown in 15 cm plates until 80% confluency was reached. GFP-Flag (Addgene 46956), wild type SF3B2-Flag, or K10R SF3B2-Flag plasmid DNA was transfected. Two days after transfection, cells were rinsed with PBS and then exposed to 500 mJ/cm^2^ UV light for crosslinking of RNA-protein in ice cold PBS. Cells were lysed in plate with ice cold RNA-IP Lysis Buffer (0.5% TritonX-100, 0.1% NP-40, 10% Glycerol, 10 mM EDTA, 20 mM Tris, 500 mM NaCl in PBS with protease inhibitor). Lysates were probe sonicated and centrifuged at 4 °C, 18,000 rcf, for 15 minutes to remove cellular debris. All samples were brought to a concentration of 10 mg/mL in 2.2 mL of RNA-IP Lysis Buffer; excess was discarded.

Input RNA was collected with Trizol (Fisher Scientific 15596018) from 200 uL of each sample. RIP was conducted from remaining lysate with Anti-Flag G1 affinity resin or Protein G Agarose (Thermo 15920010) as a negative control. IP took place for 12 hours at 4 °C and then resins were washed with ice-cold RNA-IP Wash Buffer (10% Glycerol, 10 mM EDTA, 20 mM Tris, 500 mM NaCl in PBS) before separating the resin into two aliquots.

The first aliquot was for PCR analysis of bound RNA. 100 µL of TBS, 1 µL of RNasin Ribonuclease Inhibitor (Promega N2511), and 0.5 µL of Proteinase K (New England Biolabs P8107S) was added and samples were shaken in a thermomixer at 1,200 rpm for 15 minutes at 42 °C to elute the RNA. RNA was isolated with Trizol to a final volume of 13 uL per sample. From the second resin aliquot, protein was eluted with 4% SDS and heated at 95 °C for 10 minutes. SDS-PAGE and Western blot, to assay for equal IP across samples, was conducted using 7% polyacrylamide gels and Flag antibody.

Quantitative PCR (qPCR) analysis of enriched RNA was conducted. cDNA was synthesized from the input and eluted RNA using OneScript Plus cDNA Synthesis Kit (ABM G236) and accompanying SOP and Random Primers. PCR primers targeting the previously established SF3B2 binding sites were used^18^. The negative control sample values were subtracted from the paired RIP samples to obtain true signal of specifically bound RNA. Relative pre-mRNA binding was determined by dividing values into the highest detected condition.

### AR minigene assay

The AR-v7 minigene was a gift from Dr. Xuesen Dong and we used it according to established methods^50,51^. Cells were transfected with equal amounts of the minigene and either Flag vector, WT SF3B2-Flag, or K10R SF3B2-Flag. One condition without minigene was completed as control to ensure that the observed results were due to the minigene and not endogenous RNA expression. PCR primers targeting the AR-FL or AR-v7 splice junctions were used (Table S2). Relative AR-v7/FL isoform ratio was calculated by dividing the AR-v7 splice junction expression value by the AR-FL splice junction expression value.

### qPCR

RNA was gathered using Trizol (Fisher Scientific 15596018) extraction. cDNA was synthesized from RNA using OneScript Plus cDNA Synthesis Kit (ABM G236) and accompanying SOP; Random Primers from the kit were used. qPCR was conducted with BlasTaq 2X qPCR MasterMix (abm 8G92). GAPDH was the housekeeping gene. A full primer list is in Table S2.

### Western blotting

For western blot, cells were lysed in denaturing lysis buffer unless otherwise noted and the lysate was clarified by probe sonication and centrifuge at 18,000 rcf for 15 minutes at 4 °C. Protein concentration was determined with BCA Protein Assay Kit (Thermo 23227) and 7-13% polyacrylamide gels were used for SDS-PAGE. Gels were transferred to nitrocellulose membranes (Cytiva 10600002) and blocked with either 5% bovine serum albumin (BSA) or 5% non-fat before incubation in primary antibodies diluted in 3% BSA. A full list of antibodies used in this study is in Table S3.

## Acknowledgements

This work is based on the first author’s doctoral dissertation, completed at George Washington University. The authors thank Hening Lin for his comments on this work. This work was conducted under IBC approval and was well supported by NIH grants: F31CA284578 to J. Clements, and T32CA247756 and R01CA240529-02S1 for J. Clements’s doctoral training.

## Author Contributions

J. Clements conceived the project and wrote the manuscript. J. Clements, SJ, and J. Cao completed experiments. J. Clements, LS, ACG, MG, LR, and CP created knockdown cells, fusion proteins, and cloning constructs.

## Figure Legends

**Figure S1.**
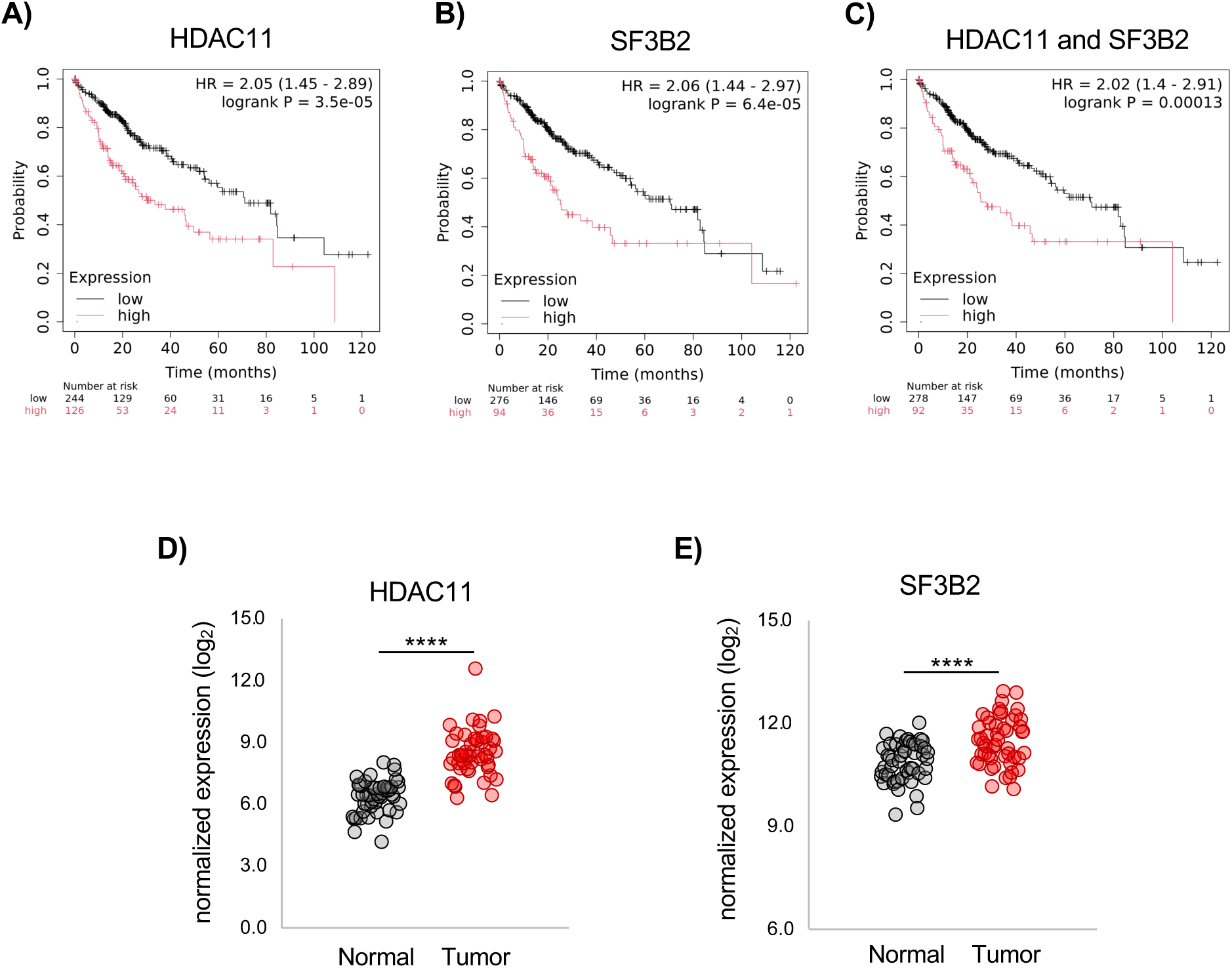
HDAC11 and SF3B2 expression in HCC patients. **(A)** HDAC11 survival plot in HCC patients**. (B)** SF3B2 survival plot in HCC patients. **(C)** Survival plot analyzing combined expression of HDAC11 and SF3B2 in HCC patients. (**D-E)** Normal versus tumor expression of HDAC11 and SF3B2 from HCC patients. Statistics are by Mann Whitney U-test. ****p<0.0001.

**Figure S2.**
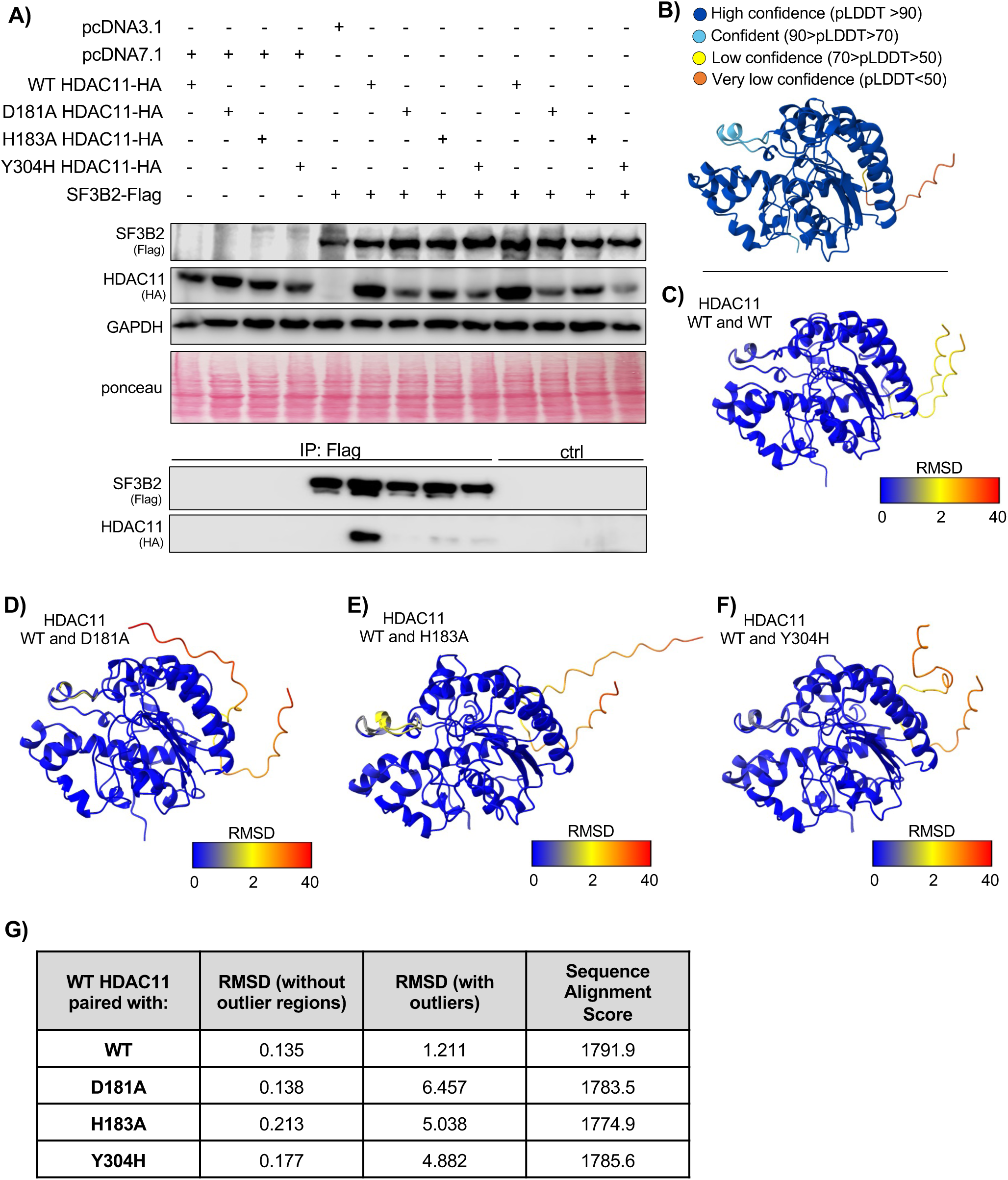
HDAC11 loss of function mutants are structurally similar to WT HDAC11 but disrupt the interaction with SF3B2. **(A)** Co-immunopurification of WT HDAC11 versus HDAC11 mutants with SF3B2**. (B)** Alpha Fold 3 predication of HDAC11 structure. **(C)** Overlay of two separate predictions of HDAC11 WT structure. **(D)** Overlay of HDAC11 WT and HDAC11 D181A structure. **(E)** Overlay of HDAC11 WT and HDAC11 H183A structure. **(F)** Overlay of HDAC11 WT and HDAC11 Y304H structure **(G)** RMSD and Sequence Alignment Scores of HDAC11 structures.

**Figure S3.**
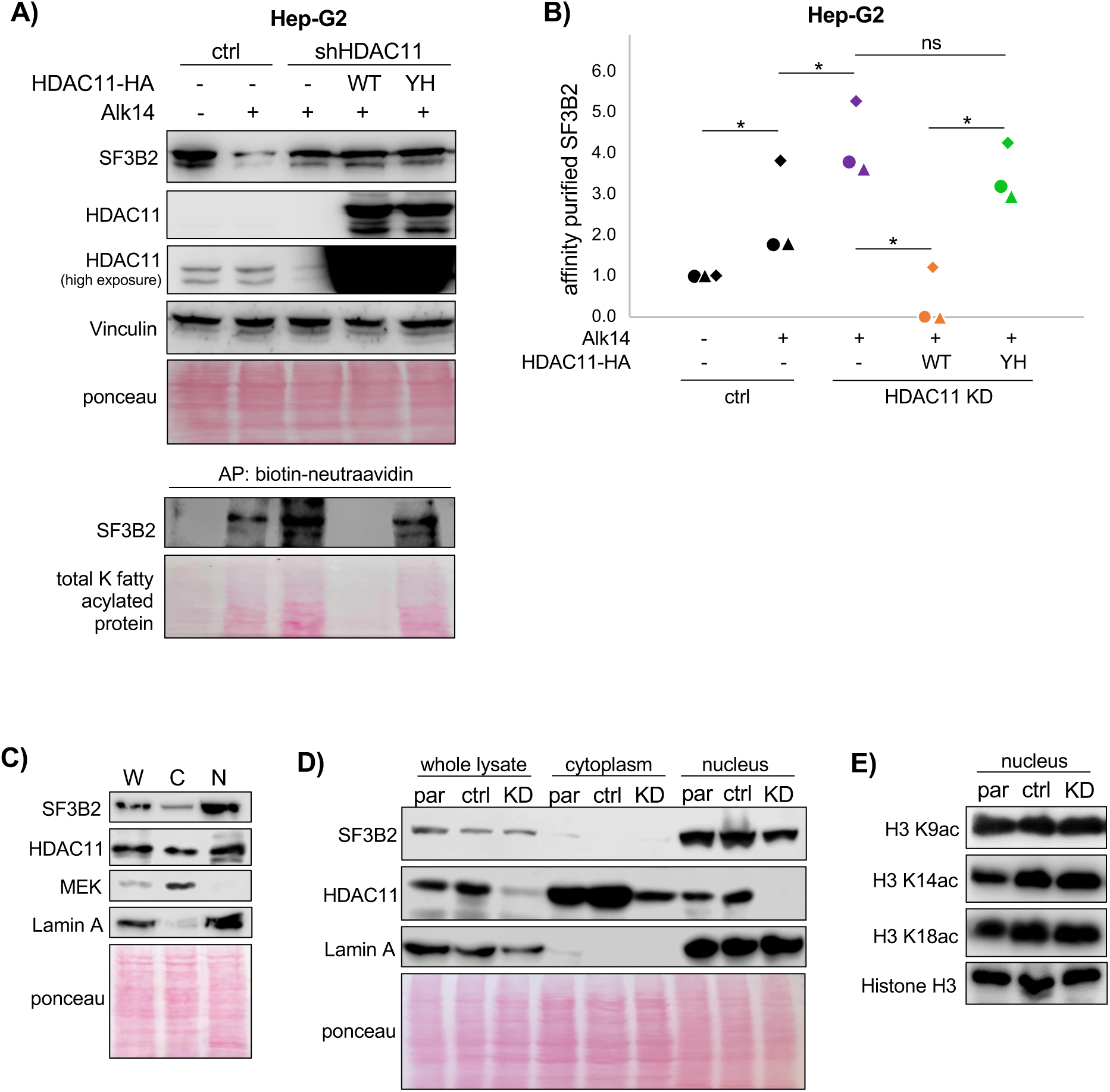
HDAC11 de-fatty acylates endogenous SF3B2 in the nucleus but does not deacetylate Histone H3. **(A)** Endogenous SF3B2 fatty acylation affinity purification in HDAC11 KD cells with overexpressed WT and Y304H HDAC11**. (B)** Quantification of H **(C).** Hep-G2 cell fractionation to assess HDAC11 and SF3B2 localization. **(D)** Localization of SF3B2 between Hep-G2 parental (par), pLKO.1 control (ctrl), or HDAC11 knockdown (KD) stable cell lines**. (E)** Western blot of H3 acetylation marks from Hep-G2 nuclear fraction.

**Figure S4.**
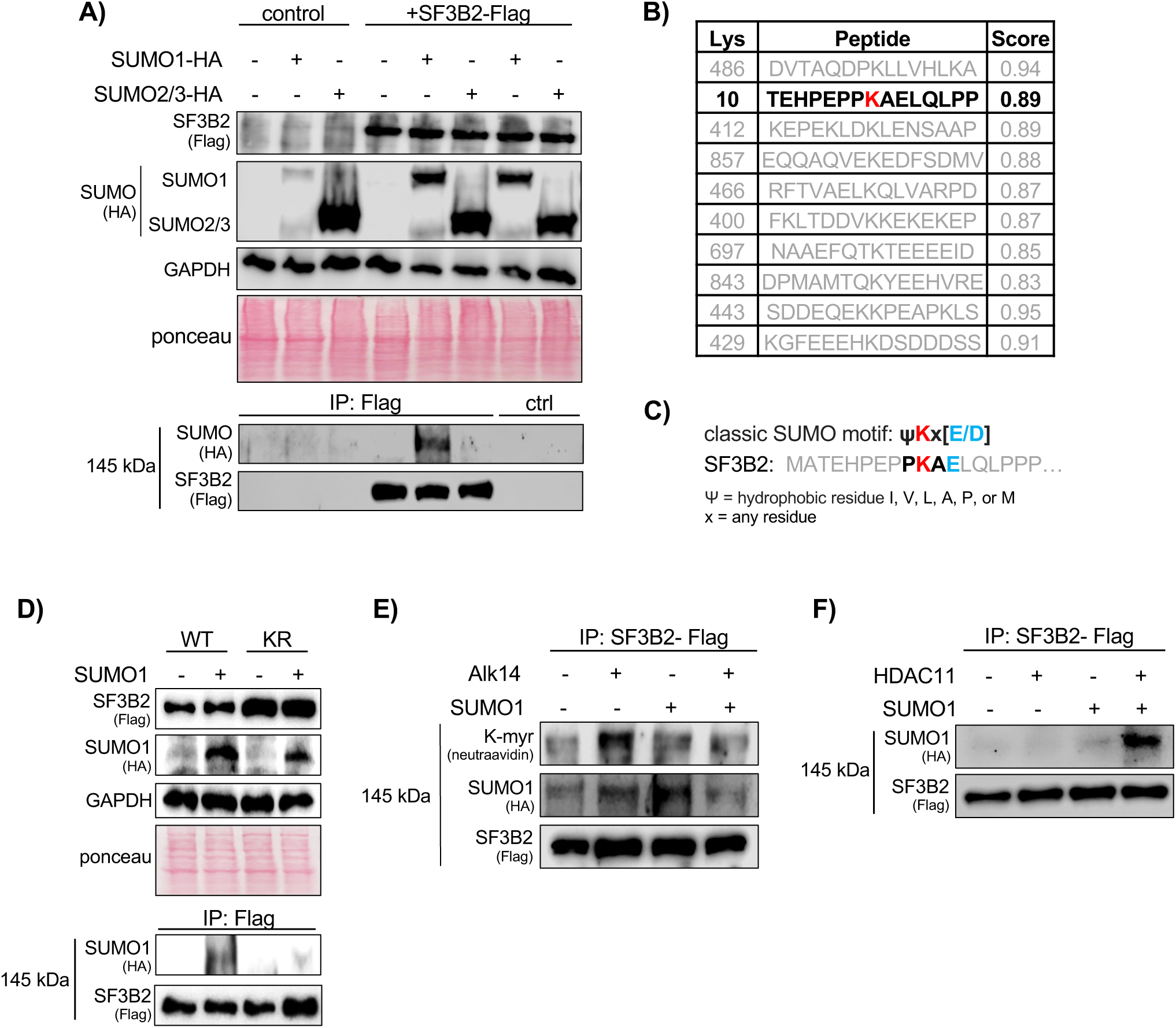
SF3B2 SUMOylation at K10 is modulated by fatty acylation. **(A)** SF3B2 can be SUMOylated by SUMO1. This is a representative image from 2 independent experiments. **(B)** Top 10 predicted SUMOylation sites on SF3B2. **(C)** K10 SUMO motif. **(D)** SF3B2 WT versus K10R SUMOylation. Image is representative of 2 independent experiments. **(E)** Analysis of SF3B2 SUMOylation versus Alk14 labelling**. (F)** Analysis of SF3B2 SUMOylation with and without HDAC11 overexpression.

**Figure S5:**
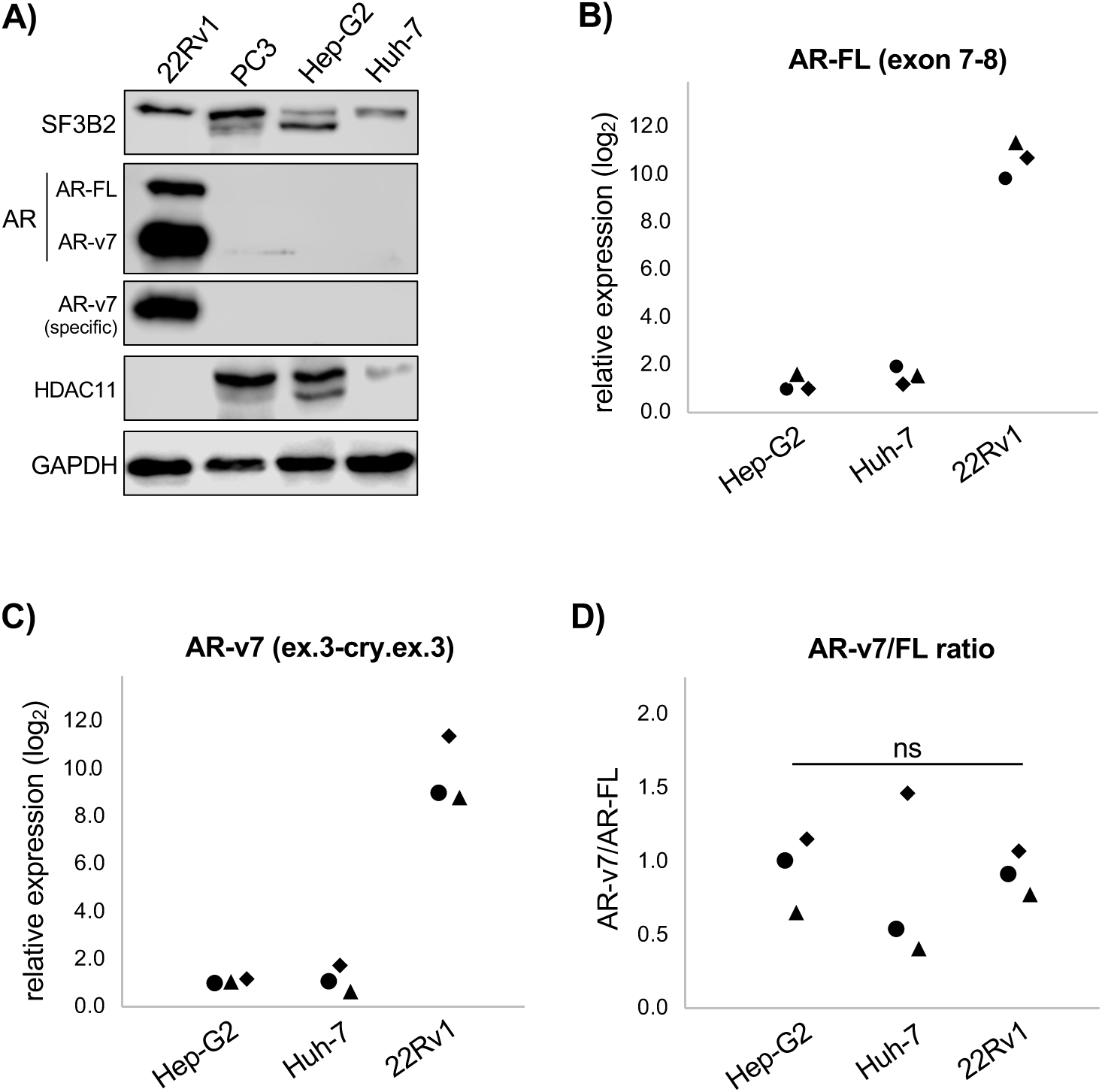
Comparison of AR-FL and AR-v7 expression in PCa and HCC cells. **(A)** Western blot comparison between 22Rv1, PC3, Hep-G2, and Huh-7 cells. PC3 cells are the negative control. **(B)** qPCR comparison of AR-FL, normalized against Hep-G2 expression level. **(C)** qPCR comparison of AR-v7, normalized against Hep-G2 expression level. **(D)** Comparison of the AR-v7/FL ratio across cell lines. Each shape in dot plots represents independent experiments. ns: not significant by ANOVA.

**Figure S6:**
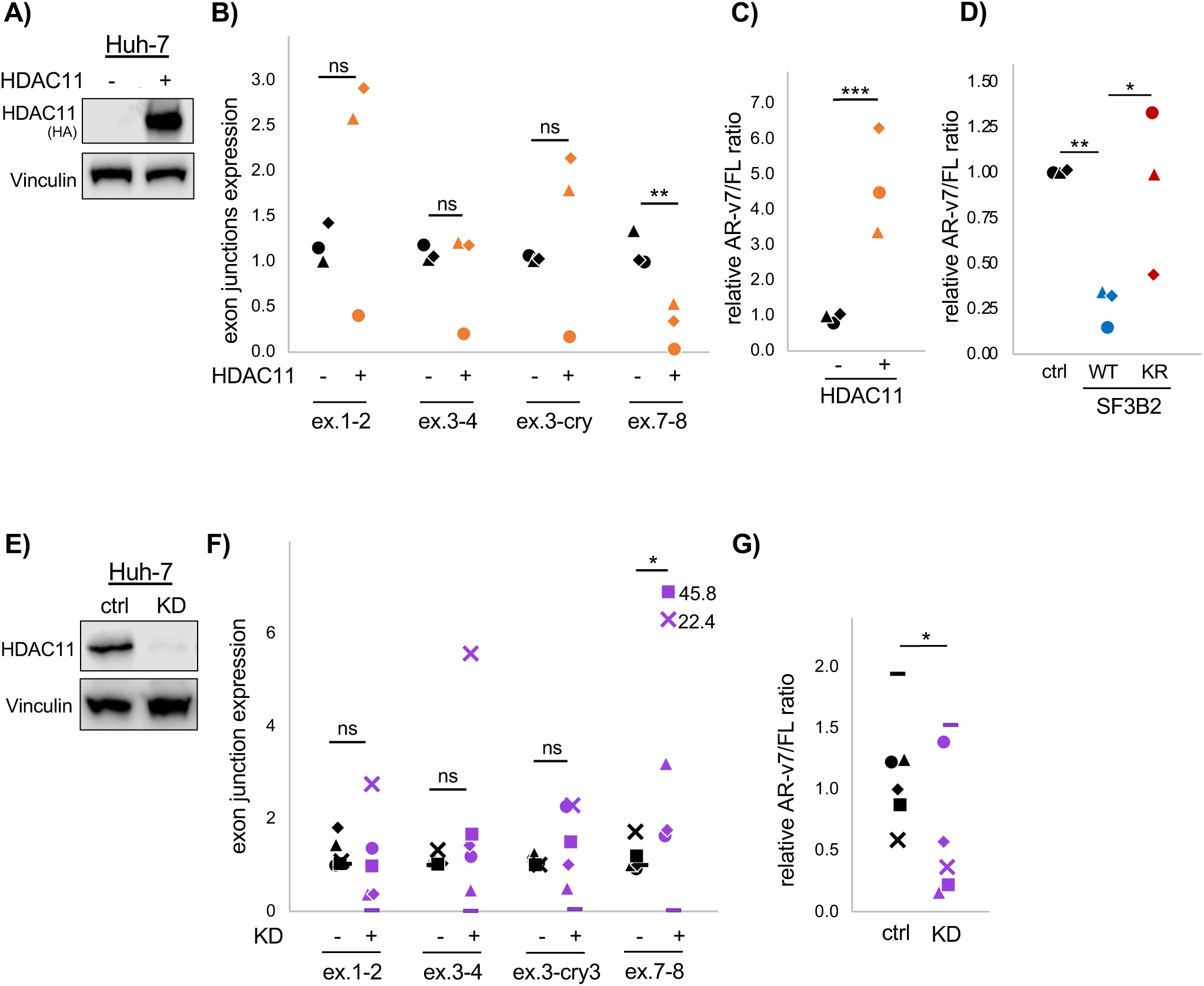
Huh-7 cells AR alternative splicing is regulated by HDAC11. **(A)** Western blot confirming overexpression of HDAC11-HA. **(B)** qPCR analysis of AR exon junctions after HDAC11 overexpression**. (C)** Relative AR-v7/FL ratio after HDAC11 overexpression**. (D)** AR-v7/FL splicing ratio with WT or K10R SF3B2**. (E)** Western blot confirming KD of HDAC11**. (F)** qPCR analysis of AR exon junctions after HDAC11 KD**. (G)** Relative AR-v7/FL ratio after HDAC11 KD. Each shape in dot plots represents independent experiments. *p<0.05, **p<0.01, ***p<0.001, ns: not significant by t-test

